# Chaperone mediated coupling of subunit availability to activation of flagellar Type III Secretion

**DOI:** 10.1101/2020.01.10.902387

**Authors:** Owain J. Bryant, Betty Y-W. Chung, Gillian M. Fraser

## Abstract

Bacterial flagellar subunits are exported across the cell membrane by the flagellar Type III Secretion System (fT3SS), powered by the proton motive force (pmf) and a specialized ATPase that enables the flagellar export gate to utilise the pmf electric potential (ΔΨ). Export gate activation is mediated by the ATPase stalk, FliJ, but how this process is regulated to prevent wasteful dissipation of pmf in the absence of subunit cargo is not known. Here, we show that FliJ activation of the export gate is regulated by flagellar export chaperones. FliJ binds unladen chaperones and, using novel chaperone variants specifically defective for FliJ binding, we show that disruption of this interaction attenuates motility and cognate subunit export. We demonstrate *in vitro* that chaperones and the FlhA export gate component compete for binding to FliJ, and show *in vivo* that unladen chaperones, which would be present in the cell when subunit levels are low, sequester FliJ to prevent activation of the export gate and attenuate subunit export. Our data indicate a mechanism whereby chaperones couple availability of subunit cargo to pmf-driven export by the fT3SS.

## Introduction

Bacterial flagella are large macromolecular rotary motors that enable cell motility. In *Salmonella,* the long helical flagellar filament propeller that extends from the cell surface is connected to a flexible hook attached to a basal body spanning the cell envelope. These flagellar substructures are assembled in a strict order, with the basal body being constructed first, followed by the hook and then the filament [1, 2]. The filament comprises thousands of copies of the major filament subunit, FliC (flagellin), together with three minor filament-class subunits: eleven copies each of FlgK and FlgL form a junction connecting the semi-rigid filament to the hook, and five FliD subunits assemble at the distal tip of the nascent flagellum to form a cap that facilitates folding and incorporation of flagellin into the growing filament [3].

Export of the filament subunits is facilitated by the cytoplasmic chaperones FlgN, FliT and FliS, which bind the C-terminal domains of their cognate structural subunits (FlgN binds FlgK and FlgL, FliT binds FliD, and FliS binds FliC) [4–6]. The chaperones then pilot their cognate subunits to the specialized flagellar Type III secretion system (fT3SS) machinery at the base of each flagellum [7–11]. Chaperone-subunit complexes initially dock at the FliI component of the flagellar export ATPase, which is evolutionarily related to the F1-ATPase [12]. Chaperoned subunits are then thought to interact with the integral membrane export gate component FlhA, where chaperones are released and the subunit cargo is unfolded for translocation across the inner membrane, *via* the FlhAB-FliPQR export gate, into a narrow central channel in the growing flagellum [9–13]. Once released from their cognate subunit cargo, the unladen chaperones FlgN and FliT - but not the flagellin chaperone, FliS - are recruited by the FliJ stalk component of the ATPase, which transfers them to newly synthesised cognate cargo to create a local cycle of chaperone-subunit binding at the membrane fT3SS [8]. This is thought to promote export of the minor filament subunits that form the hook-filament junction and the filament cap, which are required for initiation of filament assembly [14].

Export of flagellar subunit cargo across the membrane is powered by the pmf and the flagellar ATPase complex [15, 16]. Specifically, the ATPase is thought to facilitate unfolding of subunit cargo [17]. In addition, analogous to F- and V-type ATPases, ATP hydrolysis is proposed to drive rotation of the ATPase stalk, FliJ, which interacts with the nonameric export gate component, FlhA, converting the gate into a highly efficient proton-protein antiporter that utilises the ΔΨ electric component of the pmf [18]. The FliJ-FlhA interaction is essential for export gate activation, and disruption of this interaction renders the export gate inactive and attenuates protein export [18].

What is currently unclear is how FliJ activation of the export gate is regulated in the absence of subunit cargo to prevent constitutive proton influx and wasteful dissipation of the pmf. One way in which export gate activation could be regulated is by sequestration of FliJ by other binding partners, specifically the export chaperones FlgN and FliT. A role for chaperones in the regulation of T3SS activity is supported by the finding that the *Psuedomonas* virulence T3SS chaperone PcrG binds the FliJ homologue, PscO, and that loss of this interaction results in upregulation of effector secretion and renders the T3SS more resistant to pmf collapse [19]. These observations suggest that the chaperone-FliJ interaction is central to the regulation of T3SS activity, however, the mechanistic basis of this regulation remains obscure.

Here, we identify point mutant variants of the chaperones FlgN and FliT that are specifically defective for binding to the FliJ ATPase stalk yet retain the ability to bind cognate subunits and the FlhA component of the fT3SS export gate. We show *in vitro* that the FlgN and FliT chaperones compete with FlhA for binding to FliJ, and that chaperone sequestration of FliJ *in vivo* blocks the activating interaction between FliJ and the export gate. This suggests a mechanism whereby binding of FliJ by unladen chaperones acts as a signal to the export machinery that subunit cargo is unavailable and, accordingly, export activity is limited to prevent wasteful dissipation of the pmf. This model is supported by ribosome profiling data, which show that the cytoplasmic ratios of cognate subunits to chaperones are low for FlgN and FliT, relative to the ratio of flagellin to its chaperone, FliS, suggesting that levels of unladen FlgN and FliT would be a sensitive proxy measure of intracellular flagellar subunit levels. Our data indicate a mechanism by which chaperones modulate the FliJ-FlhA interaction to regulate energy use by the T3SS in response to subunit availability.

## Results

### Isolation of chaperone variants that are specifically defective for FliJ binding

To investigate the function of chaperone binding to the ATPase stalk, FliJ, we sought to identify variants of the FlgN and FliT chaperones that did not bind FliJ yet could interact with cognate subunits and the FlhA component of the fT3SS export gate. To do this, we first constructed mutant alleles of *flgN* encoding variants with three-residue deletions in the FlgN α3 helix (residues 70-102), which contains the overlapping binding sites for cognate subunits and FliJ (Fig. 1; *SI Appendix*, Fig. S1) [8]. Screening of the FlgN deletion variants using affinity chromatography pull-down assays with GST-FlgK subunit or GST-FliJ identified a single deletion variant, FlgNΔ76-78, that could bind its cognate subunit but not the ATPase stalk (Fig. 1*B*). Protein sequence comparisons revealed FlgN W_78_ to be highly conserved and replacement of this residue with alanine (FlgN-W_78_A) severely reduced FlgN binding to FliJ but did not affect binding to FlgK subunit (Fig. 1*C*; FlgNΔ90-100, which binds neither cognate subunit nor FliJ, and wild type FlgN were included as negative and positive controls, respectively) [8]. Affinity chromatography pull-down assays confirmed that FlgNΔ76-78 and FlgN-W_78_A could still bind the cytoplasmic domain of the export gate component FlhA (GST-FlhA_C_; Fig. 1*C*; interactions between FlgN variants and their binding partners are summarised in SI Appendix Table 3).

**Figure 1.**
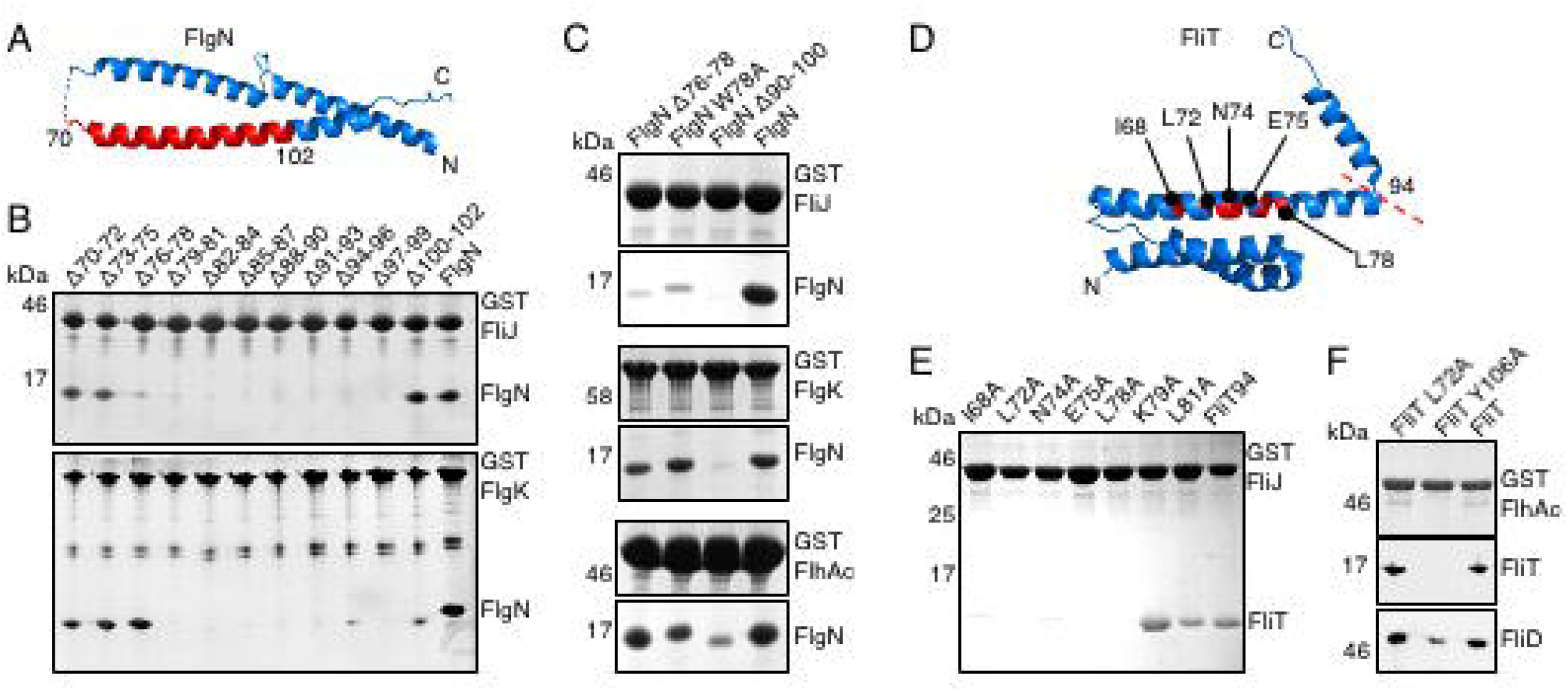
Identification of chaperone variants defective for FliJ binding. **A.** Atomic resolution structure of the 140-residue *Salmonella* FlgN (PDB: 5B3D) indicating the region (residues 70-102; red) in which three-residue scanning deletions were introduced. **B.** Affinity chromatography of glutathione sepharose-bound GST-FliJ or GST-FlgK incubated with cell lysates containing wild type FlgN or its three-amino acid deletion variants. After washing, proteins were eluted in SDS-loading buffer, separated by SDS(15%)-PAGE and stained with Coomassie Brilliant Blue. Apparent molecular weights are in kilodaltons (kDa). **C.** Affinity chromatography of glutathione sepharose-bound GST-FliJ, GST-FlgK or GST-FlhA_C_ incubated with cell lysates containing wild type FlgN or its derivative variants FlgNΔ76-78, FlgN-W78A or FlgNΔ90-100. After washing, proteins were eluted in SDS-loading buffer, separated by SDS(15%)-PAGE and stained with Coomassie Brilliant Blue. Apparent molecular weights are in kilodaltons (kDa). **D.** Atomic resolution structure of the 122-residue *Salmonella* FliT (PDB: 3A7M) indicating the position of conserved residues (red) within the region essential for FliJ binding. *In vitro* binding to FliJ is enhanced by truncating FliT at residue 94 (dashed red line). **E.** Affinity chromatography of glutathione sepharose-bound GST-FliJ with cell lysates containing FliT94 or its point mutant variants. After washing, proteins were eluted in SDS-loading buffer, separated by SDS(15%)-PAGE and stained with Coomassie Brilliant Blue. Apparent molecular weights are in kilodaltons (kDa). **F.** Affinity chromatography of glutathione sepharose-bound GST-FlhA_C_ incubated with cell lysates containing FliD and wild type FliT or its point mutant variants FliT-L72A or FliT-Y106A. After washing, proteins were eluted in SDS-loading buffer, separated by SDS(15%)-PAGE and stained with Coomassie Brilliant Blue (top panel) or were analysed by immunoblotting with anti-FliT or anti-FliD polyclonal antisera (middle and bottom panels). Apparent molecular weights are in kilodaltons (kDa).

Having identified residues in the FlgN α3 helix that, when mutated, could decouple chaperone binding of cognate subunits and FliJ, we targeted conserved residues in the equivalent α3 helix of FliT for site directed mutagenesis to alanine (Fig. 1*D*; *SI Appendix*, Fig. S1). To facilitate screening for FliJ binding, mutations were introduced into a *fliT* allele encoding a truncated variant, FliT_94_, which lacks the α4 helix and shows enhanced binding to FliJ *in vitro* [20]. Affinity chromatography pull-down assays identified five FliT_94_ variants (I_68_A, L_72_A, N_74_A, E_75_A and L_78_A) that showed severely reduced binding to GST-FliJ (Fig. 1*E*). Three of these FliJ non-binders - FliT_94_-I_68_A, FliT_94_-L_72_A and FliT_94_-N_74_A - had previously been shown to bind cognate subunit FliD [20]. For this reason, and after assessing the expression and stability of these three FliT_94_ variants in *Salmonella* (not shown), we chose to further investigate the function of truncated FliT_94_-L_72_A and full-length FliT-L_72_A.

Binding of FliT-L_72_A and truncated FliT_94_-L_72_A to cognate subunit FliD was confirmed using pull-down assays (GST-FliD; *SI Appendix*, Fig. S2). To assess binding to the export gate component FlhA, pull-down assays were carried out using GST-FlhA_C_ that had been incubated with cell lysates containing FliD cognate subunit, which is required for high affinity binding of FliT to FlhA, and either wild type FliT or full-length FliT-L_72_A (Fig. 1*F*) [9]. The FliT-Y_106_A variant, which cannot bind FlhA, was included as a negative control (Fig. 1*F*) [11]. Data from the pull-down assays indicated that, when bound to cognate subunit FliD, the FliT-L_72_A variant can bind the FlhA export gate (Fig. 1*F*; interactions between FliT variants and their binding partners are summarised in SI Appendix, Table 4).

### Attenuation of cell motility and cognate subunit export by specific disruption of the chaperone-FliJ interaction

Having identified chaperone variants (FlgNΔ76-78, FlgN-W_78_A, FliT-L_72_A) that were specifically defective for FliJ binding yet retained the ability to bind cognate subunits and the fT3SS export gate, we went on to investigate the importance of the chaperone-FliJ interaction for cell motility and subunit export. To enable expression of chaperones at physiological levels, *Salmonella* strains were engineered to carry variant chaperone genes (*flgNΔ76-78*, *flgN-W_78_A* or *fliT*-*L_72_A*) at the natural genetic loci, replacing the wild type chaperone genes. As negative controls, isogenic *Salmonella* strains in which the chaperone genes were either deleted (Δ*flgN* or Δ*fliT*) or replaced with variant genes encoding export defective chaperones unable to bind either cognate subunit (*flgNΔ90-100*) or the FlhA export gate (*fliT-Y_106_A*) were also constructed. These strains were assessed for swimming and swarming motility, and for subunit export efficiency relative to wild type *Salmonella* (Fig. 2).

**Figure 2.**
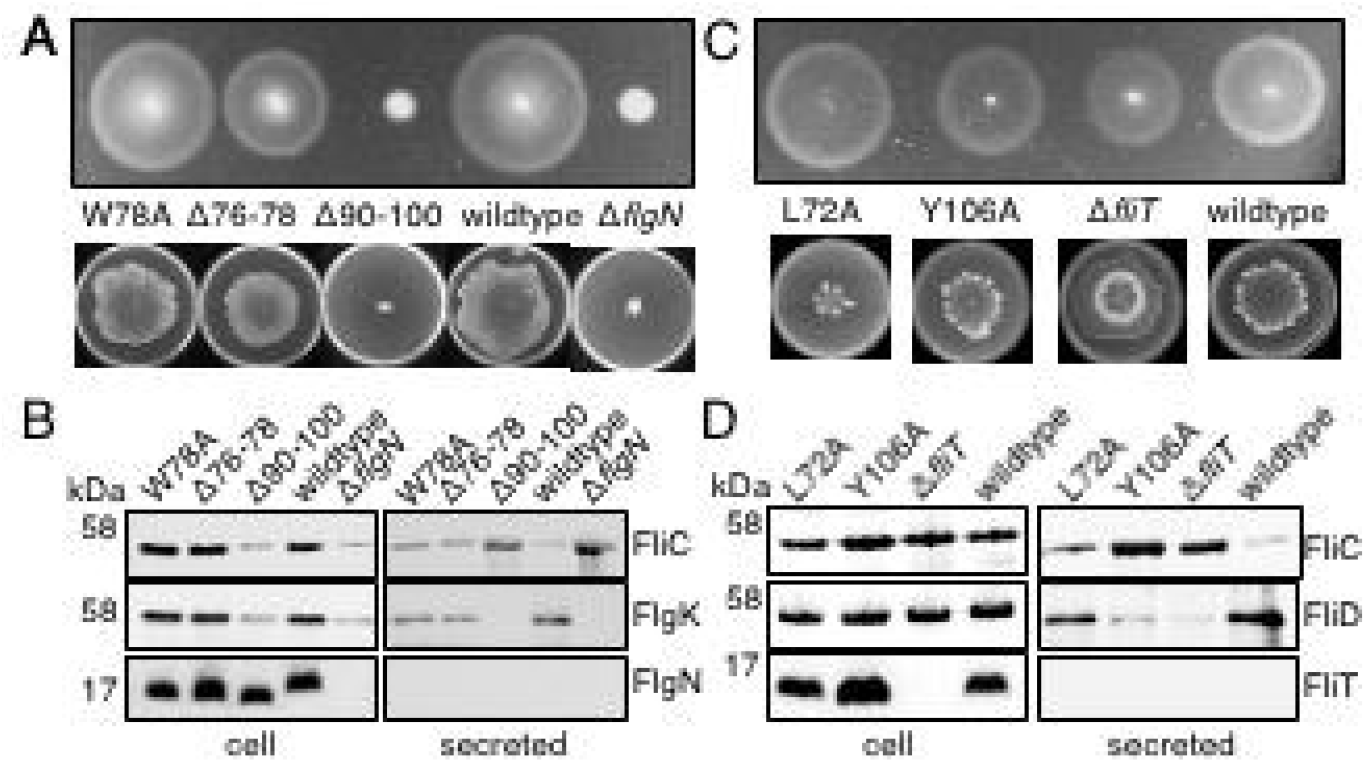
FliJ binding to FlgN and FliT is required for efficient cell motility and subunit export. **A.** Swimming (top panel; 0.25% soft tryptone agar) and swarming (bottom panels; 0.6% agar-tryptone with 0.5% glucose) motility of recombinant *Salmonella* strains producing chromosomally-encoded FlgN variants (W78A, Δ76-78, or Δ90-100), the SJW1103 wild type, and the *Salmonella flgN* null strain (Δ*flgN*). **B.** Whole cell (cell) and supernatant (secreted) proteins from late exponential-phase cultures of *Salmonella* strains producing chromosomally-encoded FlgN variants (W78A, Δ76-78, or Δ90-100), the SJW1103 wild type, and the *Salmonella flgN* null strain (Δ*flgN*) were separated by SDS (15%)-PAGE and analysed by immunoblotting with anti-FliC, anti-FlgK or anti-FlgN polyclonal antisera. Apparent molecular weights are in kilodaltons (kDa). **C.** Swimming (top panel; 0.25% soft tryptone agar) and swarming (bottom panels; 0.6% agar-tryptone with 0.5% glucose) motility of recombinant *Salmonella* strains producing chromosomally-encoded FliT variants (L72A and Y106A), the SJW1103 wild type, and the *Salmonella fliT* null strain (Δ*fliT*). **D.** Whole cell (cell) and supernatant (secreted) proteins from late exponential-phase cultures of *Salmonella* strains producing chromosomally-encoded FliT variants (L72A and Y106A), the SJW1103 wild type, and the *Salmonella fliT* null strain (Δ*fliT*) were separated by SDS (15%)-PAGE and analysed by immunoblotting with anti-FliC, anti-FliD or anti-FliT polyclonal antisera. Apparent molecular weights are in kilodaltons (kDa).

Cell populations producing the FlgNΔ76-78 or FlgN-W_78_A variants, which cannot bind FliJ, showed slightly decreased swimming (70-90%) and swarming (75-90%) motility relative to wild type *Salmonella*, though their motility was significantly better than that of the *flgN* null strain or cells producing FlgNΔ90-100, which binds neither cognate subunits nor FliJ (Fig. 2*A* and *SI Appendix*, Fig. S2). Secretion of cognate subunit FlgK was also reduced to *c.*50% of wild type in cells producing either FlgNΔ76-78 or FlgN-W_78_A, and this was accompanied by a concomitant increase in soluble extracellular FliC (to 140-150% of wild type), suggesting that assembly of the FlgKL hook-filament junction was impaired, causing inefficient initiation of filament assembly and increased release of unpolymerized FliC into culture supernatants (Fig. 2*B*) [4]. As expected, this secretion defect was more severe in the *flgN* null strain and in cells producing FlgNΔ90-100 (Fig. 2*B*).

Analysis of motility and export phenotypes associated with the FliT chaperone is complicated by the additional function of FliT as a negative regulator of the flagellar master transcriptional activator FlhD_4_C_2_ [21–23]. Cells lacking FliT show increased cellular levels of FlhD_4_C_2_, and a concomitant increase in flagellar gene expression that partially alleviates the motility and export defects typically associated with loss of chaperone activity [4]. The increased flagellar gene expression in the *fliT* null is reflected by an increase in the cellular level of non-cognate subunit FlgK, which was observed to be 1.4-fold higher than in wild type (*SI Appendix*, Fig. S3). Similarly, the level of FlgK was 1.4-fold higher in the strain producing the export defective FliT-Y_106_A (*SI Appendix*, Fig. S3), indicating increased flagellar gene expression that may be caused by sequestration of FliT-Y_106_A by cytosolic FliD, which would prevent FliT-Y_106_A binding to FlhC and targeting it for degradation [21–23]. However, cellular levels of FlgK in cells producing FliT-L_72_A were similar to wild type, suggesting that while FliT-L_72_A cannot bind FliJ, it retains the ability to bind FlhC and thus regulate flagellar gene expression (*SI Appendix*, Fig. S3).

Cells producing FliT-L_72_A displayed wild type swimming motility but showed a significant defect in population swarming motility relative to wild type *Salmonella* (*c.*50% of wild type; Fig. 2A). Moreover, export of cognate subunit FliD was reduced to *c.*50% of wild type in the FliT-L_72_A strain. These data suggest that the FliT-FliJ interaction may be required for export of sufficient FliD to promote the increased flagellation required for swarming [24, 25]. Cells producing FliT-Y_106_A, which cannot bind FlhA, showed only a slight defect in swimming and swarming motility, while the *fliT* null strain displayed a marginal reduction in swimming to *c.*80% of wild type, yet an increase in swarming motility (to *c.*110% of wild type; Fig. 2B). Both the *fliT* null and FliT-Y_106_A strain showed a significant reduction in FliD export, to 50-60% of wild type, accompanied by concomitant release of unpolymerized FliC to culture supernatants (to 110% of wild type; Fig. 2B). The confounding influence of increased flagellar gene expression in these strains, however, makes it difficult to interpret the motility and export phenotypes of the *fliT* null and FliT-Y_106_A controls, particularly as overexpression of *flhDC* is known to overcome motility defects associated with chaperone loss [26, 27].

Notwithstanding the difficulties in interpreting the motility and flagellar export phenotypes associated with loss or mutation of *fliT*, the data for FlgNΔ76-78, FlgN-W_78_A and FliT-L_72_A indicate that specific loss of chaperone interactions with FliJ, which would break the local chaperone cycle at the fT3SS, result in attenuation of cognate subunit export and cell motility. That motility and subunit export are only partially attenuated by disruption of the chaperone-FliJ interaction indicates that the variant chaperones are still able to pilot cognate subunits to the fT3SS machinery to promote export. The data suggest that chaperone binding by FliJ increases the efficiency of, but is not essential for, cognate subunit export (Fig. 2). This raises a question as to whether the chaperone-FliJ interaction might have an additional function related to the primary role of FliJ in activating the export machinery to couple efficient use of the pmf to protein export [18].

### Unladen chaperones disrupt the FliJ-FlhA_C_ interaction

The identification and characterisation of FlgN and FliT chaperone variants that retained their subunit piloting function but were specifically unable to bind the FliJ ATPase stalk opened up the possibility of further investigations into whether chaperones might regulate FliJ activation of the fT3SS. To test whether unladen chaperones could bind FliJ and block the FliJ interaction with the FlhA export gate, we developed an *in vitro* competition assay in which FlhA_C_ binding to cobalt bead-immobilized (His)_6_-FliJ was assessed in the presence of chaperone variants that could interact with FliJ but not FlhA_C_ (FliT_94_ or FlgN-Y_122_A) or were defective in binding both FliJ and FlhA_C_ (FliT_94_-L_72_A or FlgN-Y_122_AΔ76-78; Fig. 3*A*). The chaperone variants were engineered to be defective for binding to FlhA_C_ (by introduction of the FliT_94_ or FlgN-Y_122_A mutations) to prevent chaperone-FlhA_C_ interactions and thus allow specific analysis of the effect of chaperone-FliJ binding on the FlhA_C_-FliJ interaction. This assay, therefore, specifically interrogates the effect that chaperones have on the FliJ-FlhAc interaction in the absence of interfering chaperone-FlhA_C_ interactions (Fig.3A; [8–11]).

**Figure 3.**
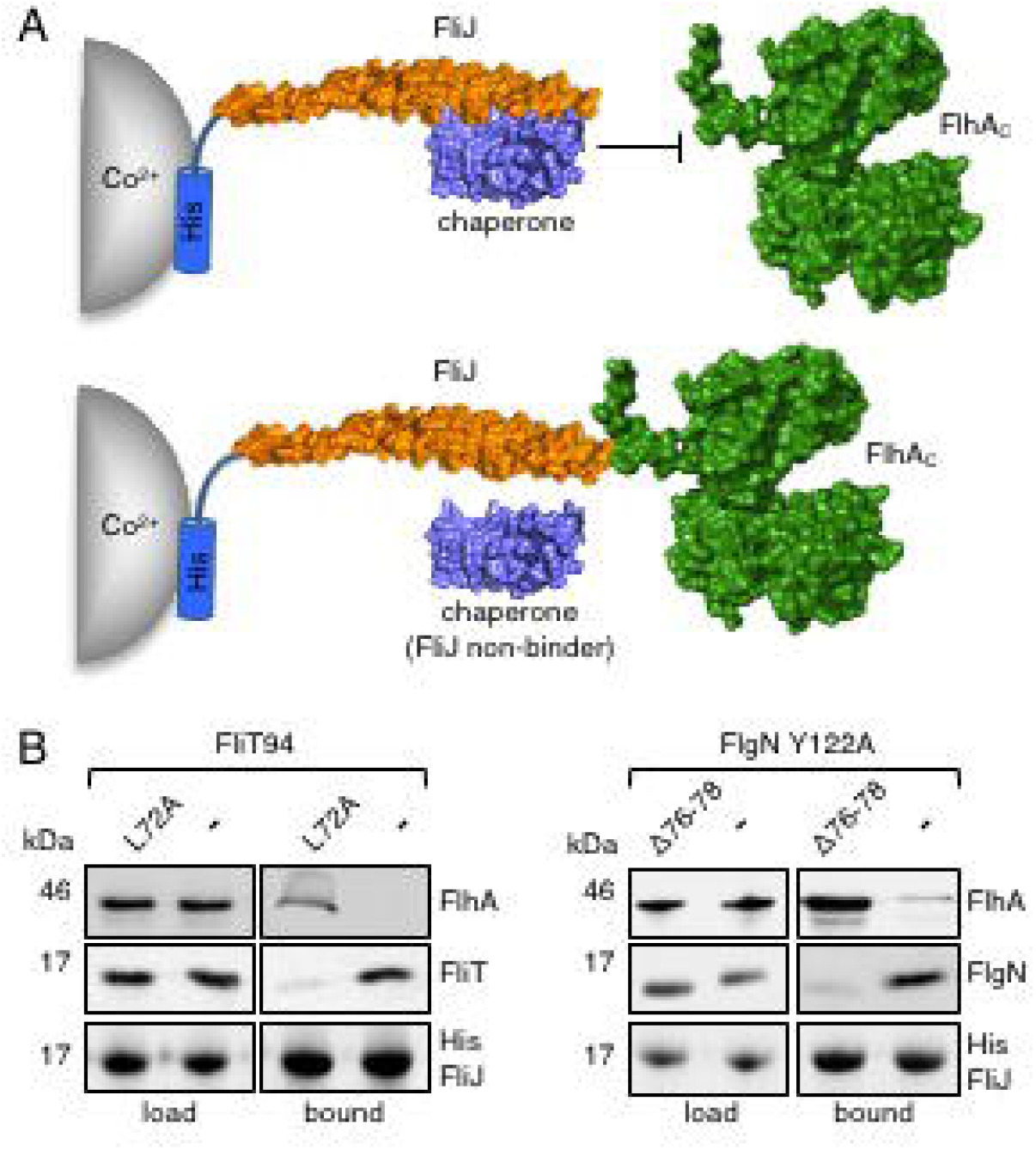
FlgN and FliT chaperones compete with FlhA_C_ for binding to FliJ *in vitro*. **A.** *In vitro* competition assay to assess FlhA_C_ binding to cobalt (Co^2+^) bead-immobilized (His)_6_-FliJ in the presence of the FlgN or FliT chaperones, or chaperone variants that cannot bind FliJ (FliJ non-binder). **B.** Cell lysates of *E.coli* C41 expressing FliT94, which cannot bind FlhA_C_ (-), or its L72A variant, which cannot bind FlhA_C_ or FliJ (left-hand panels), or FlgN Y112A which cannot bind FlhA_C_ (−) or its Δ76-78 variant, which cannot bind FlhA_C_ or FliJ (right-hand panels) were mixed with *E.coli* C41 lysates containing FlhA_C_ (load) and incubated with cobalt bead-immobilized (His)_6_-FliJ (bound). After washing, proteins were eluted from the cobalt beads in SDS-loading buffer, separated by SDS(15%)-PAGE and analysed by immunoblotting with anti-FlhA, anti-FliT, anti-FlgN or anti-FliJ polyclonal antisera. Apparent molecular weights are in kilodaltons (kDa).

As anticipated, we found that FlhA_C_ could bind (His)_6_-FliJ in the presence of chaperone variants that could not bind FliJ (Fig. 3*B*). However, when assays were performed with chaperones that could bind FliJ, the amount of FlhA_C_ pulled down by FliJ was severely reduced (Fig. 3*B*). These data show that FlgN and FliT can compete with FlhA_C_ for binding to FliJ, *i.e.* unladen chaperones bound to FliJ prevent the FliJ-FlhA_C_ interaction – an interaction known to be essential for activation of the export gate to efficiently couple pmf use to protein export [18].

### Regulation of fT3SS export gate activation in vivo by chaperone sequestration of the FliJ ATPase stalk

The *in vitro* competition assay established that the FlgN and FliT chaperones could block FliJ binding to FlhA_C_, but did not show whether activation of the FlhA_C_ export gate was prevented. To investigate this, an *in vivo* competition assay was developed in which the effect of FlgN-FliJ binding on FlhA_C_ export activity was tested under conditions where the availability of cognate subunits for FlgN was limited. Specifically, FlgN variants that could bind FliJ but not FlhA_C_ (FlgN-Y_122_A) or were defective in binding both FliJ and FlhA_C_ (FlgN-Y_122_AΔ76-78) were expressed *in trans* in a Δ*flgE* strain, in which flagellar assembly does not progress beyond completion of the rod, and the FlgN cognate subunits, FlgK and FlgL, are produced at very low levels (*SI Appendix*, Fig. S4) [28–30]. As cognate subunit levels are low in the Δ*flgE* strain, we hypothesised that *in trans* overexpression of *flgN-Y_122_A* would produce an excess of unladen FlgN-Y_122_A that would bind FliJ and disrupt the FliJ-FlhA_C_ interaction, preventing activation of the flagellar export gate and inhibiting subunit export. Conversely, we hypothesised that similar overproduction of the FlgN-Y_122_AΔ76-78 variant, which cannot bind FliJ, would not disrupt the FliJ-FlhA_C_ interaction, and would permit FliJ-dependent activation of the export gate and efficient subunit export.

Assays of subunit export in a strain producing FlgN-Y_122_A, which binds FliJ, demonstrated a *c.*75% reduction in export of the FlgD subunit compared to a strain producing FlgN-Y_122_AΔ76-78, which cannot bind FliJ (Fig *3C*). The data indicate that when unladen chaperones are present in the cell (as would occur when their cognate subunit levels are low) non-cognate subunit export is attenuated by the unladen chaperones specifically binding to FliJ, preventing export gate activation.

Taken together, the data from the *in vitro* and *in vivo* competitive binding assays indicate that unladen chaperones block the FliJ-FlhA interaction and prevent FliJ-dependent activation of the flagellar export machinery. How, then, are unladen chaperones released from FliJ, enabling export activity to be restored once cognate subunit levels return to normal? Evans and colleagues previously demonstrated that cognate subunits can capture unladen chaperones from FliJ, which would effectively free FliJ to bind the FlhA export gate and restore subunit export [8]. This provides an elegant mechanism whereby an unladen chaperone remains docked at FliJ, inhibiting pmf-driven export, until a cognate subunit is available to capture the chaperone from FliJ, allowing export activity to resume.

### Ribosome profiling reveals low cytoplasmic ratios of minor filament subunits to their cognate chaperones, FlgN and FliT

Our accumulating data suggested that, in addition to piloting subunit cargo to the fT3SS, the FlgN and FliT chaperones might be conditional regulators of fT3SS activity, which sense and respond to the availability of subunit cargo. To function effectively in the cell, this mechanism - in which unladen chaperones are a proxy measure for subunit availability - would require cytoplasmic ratios of FlgN/FliT to their cognate subunits to be low. To obtain an indication of the cytoplasmic levels of chaperones and their cognate subunits, we used RNA sequencing (RNA-Seq) and ribosome profiling (Ribo-Seq) to assess the *Salmonella* global transcript abundance and translatome, respectively, at late-log growth phase when flagellar gene expression is at its peak [31–34]. Analysis of these data confirmed that flagellar mRNAs are amongst the most abundant and highly translated in *Salmonella* (Fig. 4*A*). The ribosome profiling data indicated that the ratio of FlgN to its cognate subunits (FlgK and FlgL) was 1:3, and the ratio of FliT to its cognate subunit FliD was 1:4. These chaperone-subunit ratios are low when compared to the 1:43 ratio of FliS chaperone to its cognate subunit FliC (Fig. 4*B*), suggesting that the presence of unbound FlgN and FliT in the cell would provide a sensitive proxy measure of cytoplasmic flagellar subunit levels. Taken together with our other findings, these data strengthen the view that, when unladen, the FlgN and FliT chaperones couple subunit availability to pmf-driven export by modulating FliJ-dependent activation of the FlhA_C_ export gate.

**Figure 4.**
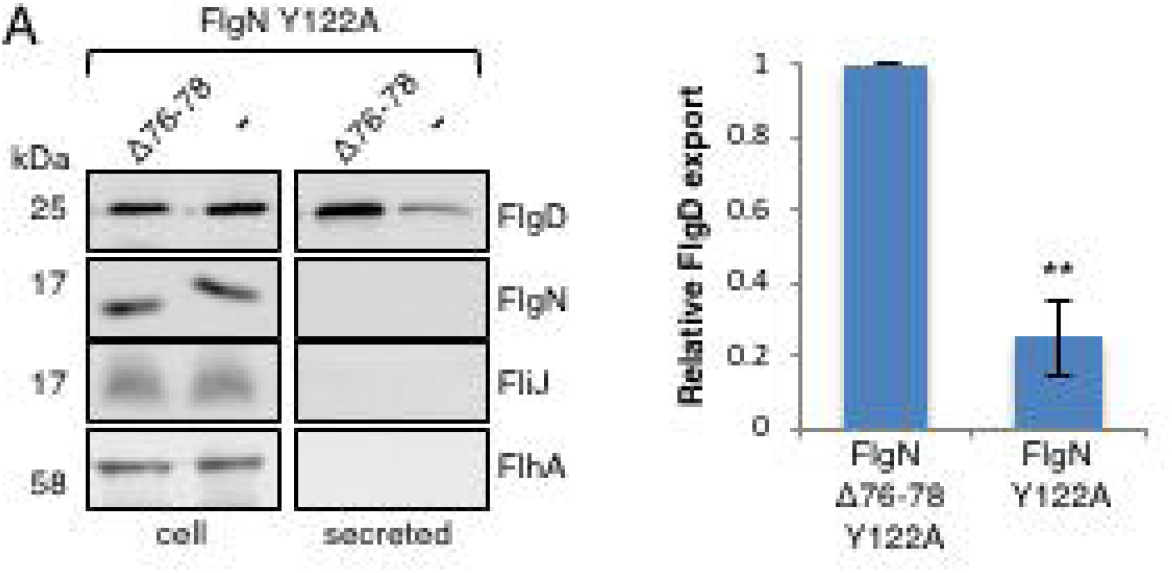
FlgN and FliT chaperones compete with FlhA_C_ for binding to FliJ *in vivo*. **A.***In vivo* competition assay to assess the effect of chaperone-FliJ binding on FlhA_C_ export activity. Export of the unchaperoned FlgD subunit was assessed in a *flgE* null strain. Whole cell (cell) and supernatant (secreted) proteins from late exponential-phase cultures of *Salmonella* Δ*flgE* strains producing pTrc99a-encoded FlgN Y122A (-) or FlgN Y122A, Δ76-78 (Δ76-78) were separated by SDS (15%)-PAGE and analysed by immunoblotting with anti-FlgD, anti-FlgN, anti-FliJ and anti-FlhA polyclonal antisera. Apparent molecular weights are in kilodaltons (kDa). Relative amounts of FlgD secreted into culture supernatants were quantified using image Studio Lite software and normalized to FlgD secreted by cells expressing the FlgN Δ76-78 Y122A variant (right-hand bar chart). Export of non-cognate subunit FlgD is attenuated in cells producing FlgN Y122A (−), which binds FliJ but not FlhA, but is significantly higher in cells producing the FlgN variant defective in binding FliJ (FlgN Δ76-78,Y122A). Data shown are the mean ± SEM of three biological replicates. *Statistically significant difference, *p* <0.01 (unpaired two-tailed *t*-test).

## Discussion

The synthesis and operation of bacterial flagella is energetically expensive, using an estimated 2% of the cell’s energy, and numerous regulatory mechanisms have evolved to increase efficiency of flagella biogenesis and function. Here, we have uncovered the mechanism by which fT3SS chaperones couple the availability of subunit cargo to the energy efficiency of flagellar export. By identifying point mutant variants of the FlgN and FliT chaperones that were specifically unable to bind the FliJ stalk component of the fT3SS ATPase, we showed that the chaperone-FliJ interaction prevents activation of the FlhA export gate component when subunit availability is low, reducing wasteful dissipation of the pmf by the fT3SS machinery.

### Mutational analysis of the FlgN and FliT chaperones reveals key residues specifically required for binding to the FliJ ATPase stalk

Our mutational analysis identified FlgN chaperone variants, FlgN W_78_A and FlgNΔ76-78, which were defective in binding to FliJ but could still pilot their cognate subunits FlgK and FlgL to dock at the export gate component FlhA_C_ (Fig.1). Motility of strains producing FlgN W_78_A or FlgNΔ76-78 was marginally reduced compared to wild type, as was export of FlgK and FlgL, indicating that loss of the FlgN-FliJ interaction reduced the efficiency of, but did not abolish, cognate subunit export (Fig. 2).

Mutational analysis of the FliT chaperone showed that several highly conserved, surface exposed residues on the helix required for cognate subunit binding were also critical for binding to FliJ (Fig.1C). Replacement of FliT leucine-72 with alanine was found to disrupt binding to FliJ, but did not affect FliT chaperone interactions with FliD cognate subunit or FlhA_C_ (Fig. 1). Cells producing FliT L_72_A did not display a swimming motility defect compared to wild type *Salmonella* (Fig. 2C). However, a marginal export defect was observed for cognate subunit FliD (Fig. 2D). Cells producing FliT L_72_A displayed reduced population swarming, indicating that the FliT-FliJ interaction may be required for the hyperflagellation associated with this form of surface motility.

### Unladen chaperones modulate the interaction of the FliJ ATPase stalk with the cytoplasmic domain of the FlhA export gate

A previous study showed that FliJ appeared to enhance the binding of a FliT/FliD complex to FlhA_C_ [9]. Using affinity chromatography of resin-bound GST-FlhA_C_ with a complex of FliT-FliD it was observed that when assays contained FliJ, binding of FliT-FliD to GST-FlhA_C_ appeared to increase [9]. The addition of excess unladen FliT chaperone to the pull-down assay reversed the FliJ-dependent binding enhancement [9]. As other studies have shown that unladen FliT chaperone does not bind FlhA_C_, we propose that FliT would block the FliJ-FlhA_C_ interaction (Figs. 4 and 5), preventing the FliJ-dependent enhancement of FliT/FliD binding to FlhA_C_, as observed by Bange and colleagues [9, 11, 35].

**Figure 5.**
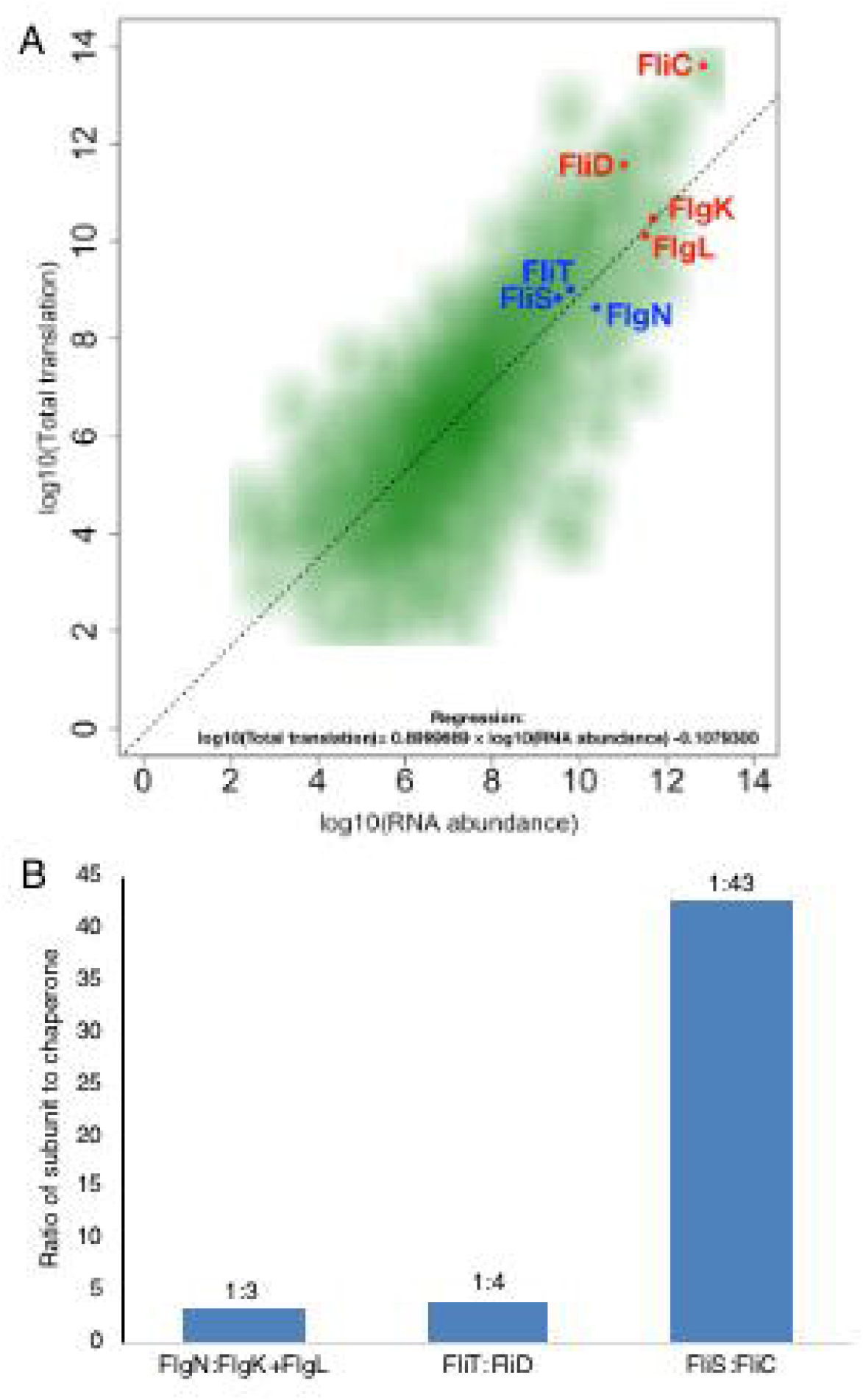
Ribosome profiling reveals cytoplasmic ratios of flagellar subunits to their cognate chaperones. **A.** Smoothed scatter plot of total translation (Ribosome profiling, measurement of ribosome occupation) and total mRNA abundance (RNA-Seq), showing flagellar **B.** chaperones FlgN, FliT and FliS (blue), their respective cognate subunits (red), and the regression between total translation and mRNA abundance (dotted line). **C.** Ratios of the normalised translation to gene length for the coding regions corresponding to chaperones and their cognate subunits (FlgN:FlgK+FlgL; FliT:FliD; FliS:FliC).

We have previously shown that FliJ-bound chaperones are efficiently captured by their cognate subunits to establish a local chaperone cycle at the membrane export machinery, which increases the efficiency of subunit export [8]. In this mechanism, the absence of cognate subunits would disrupt the chaperone cycle, resulting in unladen chaperones remaining docked at FliJ. The data presented here show that this chaperone-FliJ interaction would block FliJ-dependent activation of the flagellar export gate, preventing unnecessary activation of the export gate in the absence of subunits. In the situation where cellular levels of cognate subunit increase, chaperones would be captured from FliJ, restoring FliJ-dependent activation of the export gate and subsequent subunit export (Fig. 6).

**Figure 6.**
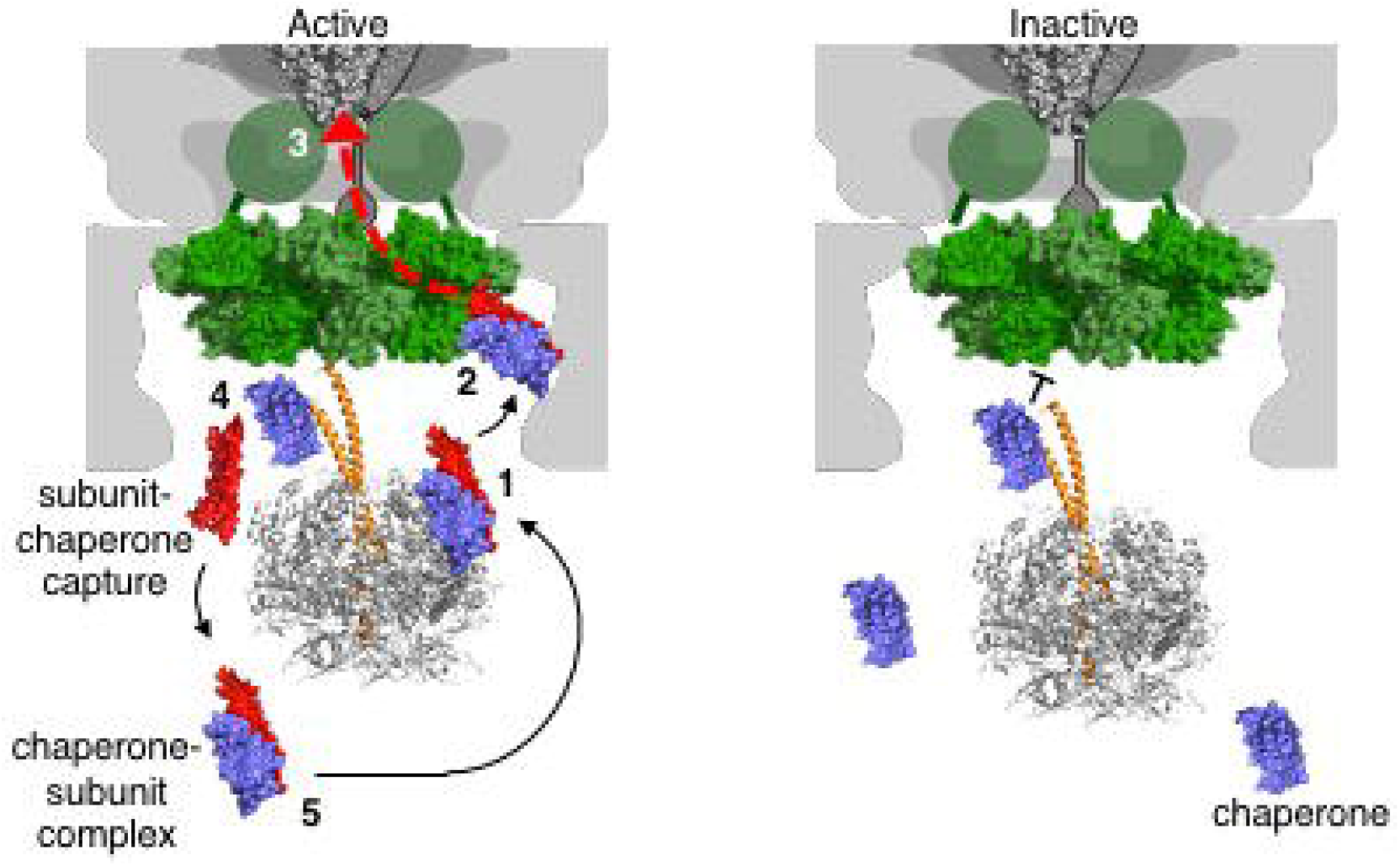
Chaperone regulation of the fT3SS machinery in response to subunit cargo availability. Chaperone-subunit complexes initially dock at the FliI component of the ATPase complex (1) before docking at the cytoplasmic domain of FlhA (2). Subunits are subsequently released from their cognate chaperones and translocated into the export channel (3). Unladen chaperones then bind FliJ from which they are captured by newly synthesized cognate subunits (4). Chaperone-subunit complexes subsequently dock at the FliI component of the ATPase, repeating the cycle (5). Under conditions where subunit availability is limited (right-hand panel), unladen FlgN and FliT chaperones remain bound to FliJ, blocking the interaction of FliJ with FlhA and reducing pmf-driven export by the fT3SS machinery.

It is well established that an interaction between FliJ and the FlhA export gate protein is required for efficient ΔΨ-driven protein export [18]. The FliH and FliI components of the ATPase support the FliJ-FlhA interaction, and hydrolysis of ATP by the ATPase complex is proposed to drive rotation of FliJ, allowing it to contact sequentially all nine monomers of the FlhA nonameric ring, converting the export machinery into a highly efficient proton-protein antiporter [12,15,16,18]. The fT3SS ATPase components share many features of the F_1_ ATP synthase: FliI and FliJ share considerable structural similarity to the α/β- and γ-subunits, respectively, and FliH shares considerable amino acid sequence similarity to the b and δ subunits [36]. Furthermore, analogous to the F_0_ ATP synthase component, the flagellar export machinery harnesses the power of the pmf [12, 37]. Multiple mechanisms have been identified that regulate both ATPase activity and pmf use by the F_1_F_0_-ATP synthase, including proteins that inhibit efficient coupling between the F_1_ and F_0_ components [37–39]. Our data demonstrate that unladen export chaperones disrupt FliJ-dependent activation of the flagellar export gate, which would prevent wasteful pmf use in the absence of cognate subunits, by effectively preventing a FliJ-FlhA interaction until the chaperone is captured from FliJ by a cognate subunit, relieving the inactive state.

### A mechanism coupling export gate activation to availability of subunits for export

Our data support a model whereby, when subunits are at low levels in the cell, unladen FlgN and FliT bind to FliJ, preventing the FliJ-FlhA interaction and subsequent activation of the export gate. Ribosome profiling revealed the amounts of chaperone and cognate subunit(s) produced by the cell and while all three flagellar chaperones are produced at similar levels (Fig. 6), the intracellular ratio of chaperone to subunit is low for the FlgN and FliT chaperones (1:3 and 1:4, respectively), compared to the ratio of FliS chaperone to flagellin (1:43), which is produced and exported at far higher levels than FlgK and FlgL [8]. As a result, the presence of unladen FlgN and FliT would serve as a sensitive proxy measure for changes in the availability of newly synthesised flagellar subunits in the cell, as these chaperones are more likely to be free, compared to the FliS chaperone, when subunit levels drop. This may in part explain why FliS has not evolved to bind FliJ.

Whether such a mechanism of chaperone mediated coupling of subunit availability to export gate activation occurs in the related vT3SS is not clear. However, a subset of export chaperones, specifically for the virulence needle translocon and tip subunits, bind to vT3SS FliJ homologues [19, 40, 41, 46]. Notably, an export chaperone (PcrG) in *Pseudomonas aeruginosa* binds to the FliJ homologue (PscO) and modulates pmf use by the vT3SS, although the mechanistic basis for this was unclear [19]. Independent to PscO (FliJ) binding, and in the absence of cognate subunit, PcrG can regulate effector secretion through interactions with the FlhA homologue, PcrD, preventing effectors accessing the export machinery [19]. These findings indicate that interactions between a subset of export chaperones and FliJ homologues may be conserved among vT3SSs. Whether these binding events also serve to regulate export gate activation with respect to subunit availability is yet to be determined.

Chaperone modulation of fT3SS activity may be important when *Salmonella* encounters environmental conditions in which flagellar gene expression is down-regulated, including during the early stages of host infection [42–45]. Our data support a model for a local chaperone cycle at the T3SS machinery that (i) facilitates efficient docking and export of cognate subunit cargo when subunit/effector availability is high, and (ii) prevents constitutive proton influx and wasteful dissipation of the pmf when subunit cargo availability is low (Fig. 6).

## Materials and Methods

### Bacterial strains, plasmids and growth conditions

Wild type *Salmonella enterica* serovar Typhimurium (*S. typhimurium*) SJW1103 is motile [48]. The Δ*fliT*∷K_m_^R^ mutant in which the *fliT* gene was replaced by a kanamycin resistance cassette was constructed using the λ Red recombinase system [49]. Strains containing chromosomally encoded FliT or FlgN variants were constructed by *aph*-I-SceI Kanamycin resistance cassette replacement using pWRG730 [49]. Bacteria were cultured at 30–37°C in Luria-Bertani (LB) broth containing, where appropriate, ampicillin (100 μg/ml) or chloramphenicol (20 μg/ml) and collected by centrifugation (6,000*g* for 10 min). Recombinant proteins were expressed in *Salmonella* from the isopropyl β-D-thiogalactoside-inducible (IPTG)-inducible plasmid pTrc99a [50].

Recombinant proteins for purification were expressed in *E.coli* C41 [51] from (IPTG)-inducible pACT7 [52] or pGEX-4T3 [53]. To construct recombinant plasmids encoding wild type or derivative genes, *Salmonella* genes were amplified from chromosomal DNA by PCR or overlap-extension PCR using Q5 High-Fidelity DNA polymerase. PCR products were inserted BamHI/XhoI into pGEX-4T3 or NdeI/BamHI into pACT7 and pTrc99a. Inserts were verified by DNA sequencing (Department of Biochemistry, University of Cambridge). A full list and description of strains and plasmids used in this study can be found in *SI Appendix*, Tables S1 and S2.

### Affinity chromatography co-purification assays

Co-purification of proteins expressed in *E.coli* C41 was performed with glutathione-sepharose 4B. Cells were resuspended in buffer A (50 mM sodium phosphate pH7.4, 150 mM NaCl, 1mM β–mercaptoethanol) and mechanically lysed with a Constant Systems cell disruptor at 30,000 kPa. Cleared lysates containing GST-tagged proteins were incubated with affinity resin for 1 hour. After extensive washing with buffer A, cell lysates containing overexpressed untagged prey proteins were then incubated with the resin pre-bound GST-bait. Resins were washed extensively with buffer A and proteins eluted by boiling in SDS-PAGE loading buffer. Proteins were separated by SDS-PAGE and immunoblotted.

### Flagellar subunit export assay

*Salmonella* strains were cultured at 37 °C in LB broth containing ampicillin and 100 μM IPTG to mid-log phase (A_600_ 0.6-0.8). Cells were centrifuged (6000*g*, 5 min), resuspended in fresh media and grown for a further 60 min at 37 °C. Cells were pelleted by centrifugation (16,000*g*, 5 min) and the supernatant passed through a 0.2 μm nitrocellulose filter. Supernatant proteins were precipitated with 10% trichloroacetic acid (TCA) and 1% Triton-X100 on ice for 1 hour, pelleted by centrifugation (16,000*g*, 10 min), washed with ice-cold acetone and resuspended in SDS-PAGE loading buffer (volumes calibrated according to cell densities). Fractions were analysed by immunoblotting.

### Motility assays

For swimming motility, *Salmonella* strains were grown in LB broth to A_600_ 1. Two microliters of culture were inoculated into soft tryptone agar (0.25% agar, 10 g/L tryptone, 5g/L NaCl). Plates were incubated at 37 °C for between 4 and 6 hours. For swarming motility, one microliter of overnight cultures grown in LB broth was inoculated onto dried tryptone agar plates (0.6% agar, 10g/L tryptone, 5g/L NaCl) supplemented with 0.3% glucose and incubated at 30 °C for 16 hours.

### Chaperone-FliJ *in vitro* competition assay

Recombinant proteins were expressed in *E.coli* C41 and cells were mechanically lysed as described for the co-purification assays. Cleared lysates containing His-tagged FliJ were incubated with cobalt affinity resin for 1 hour. After extensive washing, cell lysates containing overexpressed untagged FlhA_C_ was added to each resin fraction and an equal volume of wild type FliT_94_, FliT_94_ L_72_A, wild type FlgN or FlgN W_78_A was added. Proteins were incubated for 1 hour, the resin was washed extensively with buffer A and proteins were eluted by boiling in SDS-PAGE loading buffer. Proteins were separated by SDS-PAGE and immunoblotted.

### FlgN-FliJ *in vivo* competition assay

*Salmonella flgE* null strains carrying either pTrc99a encoding *flgN-*Y_122_AΔ76-78 or *flgN-*Y_122_A were grown in LB broth containing ampicillin and inducing agent (50 μM IPTG) to A_600_ 1.0. Cells were centrifuged (6000*g*, 5 min) and resuspended in fresh media and grown for a further 60 min at 37°C. Cells were pelleted by centrifugation (16,000*g*, 5 min) and the supernatant passed through a 0.22 μm nitrocellulose filter (Sartorius). Proteins were precipitated with 10% trichloroacetic acid (TCA) and 1% Triton-X100 on ice for 1 hour, pelleted by centrifugation (16,000*g*, 10 min, 4°C), washed with ice-cold acetone and resuspended in SDS-PAGE loading buffer (volumes calibrated according to cell densities). Fractions were analysed by immunoblotting.

### Ribosome profiling and RNA-Seq

Libraries were prepared essentially as described previously [54, 55] with the following modifications: Cells were grown in LB at 37°C with 180 RPM shaking to A_600_ 1.0. Chloramphenicol was added to the culture to a final concentration of 1500 μg/ml, followed by rapid cooling of the culture and harvesting of cells by centrifugation (6,000 g, 1 min, 4°C). The cell pellet was quickly resuspended in 1 ml ice cold bacterial profiling buffer (20 mM Tris-Cl pH 7.5, 140 mM KCl, 5 mM MgCl_2_, 1500 μg/ml chloramphenicol, 0.5 mM dithiothreitol (DTT), 0.5% NP40, 1% Triton X-100, 2.5% Sucrose) and flash-frozen for cryo-grinding in liquid nitrogen. The frozen powder was thawed and clarified by centrifugation (13,000 g, 2 mins, 4°C) followed by adjustment of A_254 nm_10. Lysates were either snap frozen in liquid nitrogen for storage at −80°C or nuclease treated with RNase 1 (700U, Ambion) followed by ribosomal RNA depletion using a bacterial ribozero kit (Illumina) prior to library preparation for ribosome profiling. For parallel RNA-Seq, total RNA was extracted from corresponding lysates followed ribosomal RNA depletion using the bacterial ribozero kit (Illumina) prior to library preparation. For Ribosome profiling, rRNA was further depleted with duplex-specific nuclease as described previously [54]. Ribosome profiling and RNA-Seq libraries were pooled and sequenced using the NextSeq® 500/550 platform and data were trimmed and mapped to the transcriptome assembly of *Salmonella enterica* subsp. *enterica* serovar Typhimurium strain ST4/74 (GenBank accession CP002487.1). Paired ribosome profiling and RNA-Seq analysis was performed with riboSeqR as previously described [55, 56].

### Quantification and statistical analysis

Experiments were performed at least three times. Immunoblot data were quantified using Image Studio Lite. The unpaired two-tailed Student’s *t*-test was used to determine *p*-values, and significance was defined as \**p* < 0.05. Data are represented as mean ± standard error of the mean (SEM), unless otherwise specified and reported as biological replicates.

## Abbreviations

T3SS: Type III secretion system
fT3SS: flagellar Type III secretion system
vT3SS: virulenceType III secretion system
pmf: proton motive force

## Acknowledgements

We thank Colin Hughes and Lewis Evans for useful discussions, and Sangita Ahmed for technical assistance. This work was funded by grants from the Biotechnology and Biological Sciences Research Council (BB/M007197/1) to G.M.F, Medical Research Council [MR/R021821/1] to B.Y-W.C., and a University of Cambridge John Lucas Walker Studentship to O.J.B.

## Footnotes

The authors declare no conflict of interest.

## Author contributions

**Owain J. Bryant**: Conceptualization, Data curation, Investigation, Formal analysis, Methodology, Visualisation, Writing – original draft, Writing – review & editing. **Betty Y-W Chung**: Data curation, Formal analysis, Funding acquisition, Methodology, Investigation, Visualisation, Software, Writing – original draft, Writing – review & editing. **Gillian M. Fraser**: Conceptualization, Data curation, Formal analysis, Funding acquisition, Investigation, Methodology, Project administration, Resources, Supervision, Visualisation, Writing – original draft, Writing – review & editing.

## Supplementary Figure Legends

**Figure S1.**

**A.** Affinity chromatography of glutathione sepharose-bound glutathione-S-transferase (GST) incubated with cell lysates containing FliT94. After washing, proteins were eluted in SDS-loading buffer, separated by SDS(15%)-PAGE and stained with Coomassie Brilliant Blue (top panel). The cell lysate load was analyzed

**B.** by immunoblotting with polyclonal anti-FliT anti-sera (bottom panel). Apparent molecular weights are in kilodaltons (kDa).

**C.** Affinity chromatography of glutathione sepharose-bound glutathione-S-transferase (GST) incubated with cell lysates containing FlgN. After washing, proteins were eluted in SDS-loading buffer, separated by SDS(15%)-PAGE and stained with Coomassie Brilliant Blue (top panel). The cell lysate load was analyzed by immunoblotting with polyclonal anti-FlgN anti-sera (bottom panel). Apparent molecular weights are in kilodaltons (kDa).

**Figure S2.**

**A.** Affinity chromatography of glutathione sepharose-bound GST-FlgL (top two panels) or glutathione-S-transferase (GST; bottom panels) incubated with cell lysates containing FlgN or its derivative variants FlgNΔ76-78, FlgN-W78A or FlgNΔ90-100. After washing, proteins were eluted in SDS-loading buffer, separated by SDS(15%)-PAGE and stained with Coomassie Brilliant Blue to visualise GST-FlgL and GST, or analysed by immunoblotting with polyclonal anti-FlgN anti-sera. The cell lysate load (FlgN load) was also analyzed by immunoblotting with polyclonal anti-FlgN anti-sera. Apparent molecular weights are in kilodaltons (kDa).

**B.** Affinity chromatography of glutathione sepharose-bound GST-FliJ (top two panels), GST-FliD (middle two panels) or glutathione-S-transferase (GST; bottom panels) with cell lysates containing FliT and its mutant variants. After washing, proteins were eluted in SDS-loading buffer, separated by SDS(15%)-PAGE and stained with Coomassie Brilliant Blue to visualise GST-FliJ, GST-FliD and GST, or analysed by immunoblotting with polyclonal anti-FliT anti-sera. The cell lysate load (FliT load) was also analyzed by immunoblotting with polyclonal anti-FliT anti-sera. Apparent molecular weights are in kilodaltons (kDa).

**C.** Cell lysates of *E.coli* C41 expressing FliT (top panel) or FliD (bottom panel) used as load fractions in affinity chromatography assays with GST-FlhA_C_ in Figure 1. Proteins were separated by SDS(15%)-PAGE and analysed by immunoblotting with polyclonal anti-FliT anti-sera or polyclonal anti-FliD anti-sera.

**Figure S3.**

**A**. Whole cell and supernatant proteins from late exponential-phase cultures of *Salmonella* strains producing chromosomally-encoded FliT variants (L72A and Y106A), the *Salmonella fliT* null strain (Δ*fliT*), and the SJW1103 wild type were separated by SDS (15%)-PAGE and analysed by immunoblotting with anti-FlgK polyclonal antisera. Apparent molecular weights are in kilodaltons (kDa).

**Figure S4.**

**A.**Whole cell proteins from late exponential-phase cultures of a *Salmonella flgD* null strain (Δ*flgD*), and the SJW1103 wild type were separated by SDS (15%)-PAGE and analysed by immunoblotting with anti-FlgK polyclonal antisera. Apparent molecular weights are in kilodaltons (kDa).

