## Supplementary figures and images for "Chaperone mediated coupling of subunit availability to activation of flagellar Type III Secretion"

### Supplementary Figures

*Supplementary Information (SI)*
*Appendix 1*

#### Figure S1

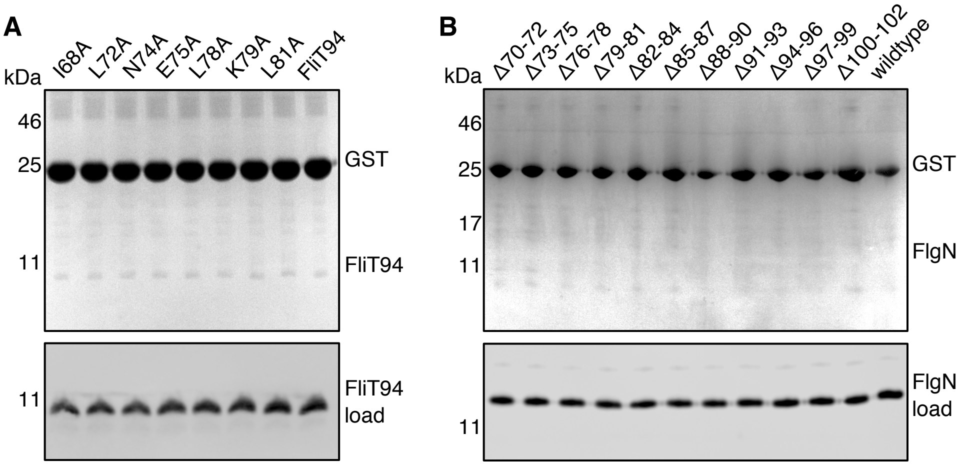

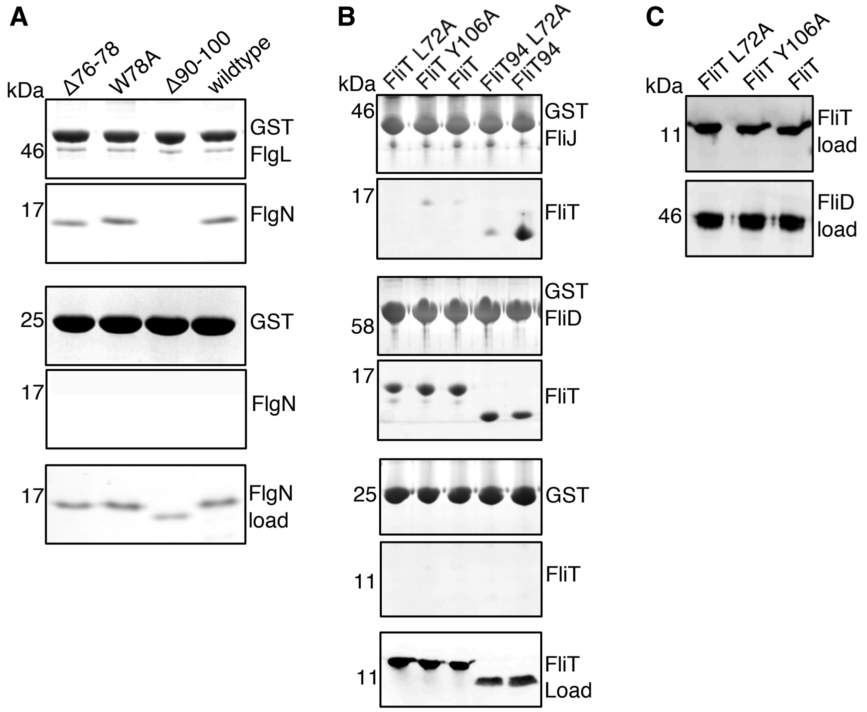

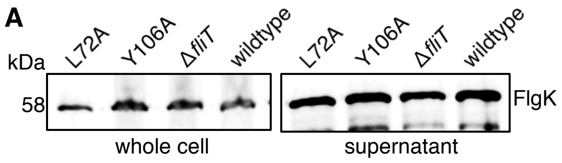

#### Figure S4

**A**

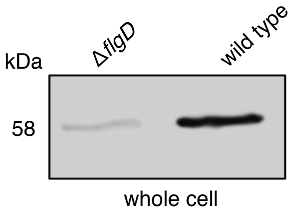
