## Supplementary Tables for "Chaperone mediated coupling of subunit availability to activation of flagellar Type III Secretion"

### Supplementary Table 1

| **Strains** | **Description** |
|---|---|
| ***Salmonella enterica serovar Typhimurium (S. typhimurium)*** | |
| SJW1103 | wildtype |
| *fliT* null | Δ*fliT*::kmR |
| *fliT* L72A | *fliT* Leu72Ala |
| *fliT* Y106A | *fliT* Tyr106Ala |
| *flgN* null | Δ*flgN*::kmR |
| *flgN* W78A | *flgN*Trp78Ala |
| *flgN* Δ76-78 | FlgN1-75, 79-140aa |
| *flgN* Δ90-100 | FlgN1-89, 101-140aa |
| ***Escherichia coli*** | |
| C41 | *F ompT gal dcm* $hsdS_B$ $(r_B^- \ m_B^-)$*(44)* |

### Supplementary Table 2

| | |
|---|---|
| **Plasmids** | |
| pGEX-4T3 FliJ | GST fusion, 1-147aa |
| pGEX-4T3 $FlhA_C$ | GST fusion, 328-692aa |
| pGEX-4T3 FliD | GST fusion, 1-467aa |
| pGEX-4T3 FliI | GST fusion, 1-456aa |
| pGEX-4T3 FlgK | GST fusion, 1-553aa |
| pGEX-4T3 FlgL | GST fusion, 1-317aa |
| pACT7 FliT94 | 1-94aa |
| pACT7 FliT94 I60A | 1-94aa, Iso60Ala |
| pACT7 Fli94 D62A | 1-94aa, Asp62Ala |
| pACT7 FliT94 V64A | 1-94aa, Val64Ala |
| pACT7 FliT94 Y67A | 1-94aa, Tyr67Ala |
| pACT7 FliT94 I68A | 1-94aa, Iso68Ala |
| pACT7 FliT94 L72A | 1-94aa, Leu72Ala |
| pACT7 FliT94 N74A | 1-94aa, Asn74Ala |
| pACT7 FliT94 E75A | 1-94aa, Glu75Ala |
| pACT7 Fli94 L78A | 1-94aa, Leu78Ala |
| pACT7 FliT94 K79A | 1-94aa, Lys79Ala |
| pACT7 FliT94 L81A | 1-94aa, Leu81Ala |
| pACT7 FliT | 1-122aa |
| pACT7 FliT L72A | 1-122aa, Leu72Ala |
| pACT7 FliT Y106A | 1-122aa, Tyr106Ala |
| pACT7 FlgN Y122A | 1-140aa, Tyr122Ala |
| pACT7 FlgN Y122A, Δ76-78 | 1-75, 79-140aa, Tyr122Ala |
| pACT7 FlgN Δ70-72 | 1-69, 73-140aa |
| pACT7 FlgN Δ73-75 | 1-72, 76-140aa |
| pACT7 FlgN Δ76-78 | 1-75, 19-140aa |
| pACT7 FlgN Δ79-81 | 1-78, 82-140aa |
| pACT7 FlgN Δ82-84 | 1-81, 85-140aa |
| pACT7 FlgN Δ85-87 | 1-84, 88-140aa |
| pACT7 FlgN Δ88-90 | 1-87, 91-140aa |
| pACT7 FlgN Δ91-93 | 1-90, 94-140aa |
| pACT7 FlgN Δ94-96 | 1-93, 97-140aa |
| pACT7 FlgN Δ97-99 | 1-96, 100-140aa |
| pACT7 FlgN Δ100-102 | 1-99, 103-140aa |
| pACT7 FlgN W78A | 1-140aa, Trp78Ala |
| pACT7 FlgN Δ90-100 | 1-89, 101-140aa |
| pACT7 FlgN | 1-140aa |
| pET15b FliJ | 1-147aa, Nt-Hisx6 |
| pACT7 $FlhA_C$ | 328-692aa |
| pTrc99a FlgN Y122A | 1-140aa, Tyr122Ala |
| pTrc99a FlgN Y122A, Δ76-78 | 1-75, 79-140aa, Tyr122Ala |

### Supplementary Table 3

Table summarising interactions between FlgN variants (left column) and export machinery components or cognate subunits (top row). ✓ refers to binding and x refers to weak or no binding.

| **Protein** | FliJ | FlhA | FlgK | FlgL |
|---|---|---|---|---|
| FlgNΔ76-78 | x | ✓ | ✓ | ✓ |
| FlgN Y122A | ✓ | x | ✓ | ✓ |
| FlgNΔ76-78, Y122A | x | x | ✓ | ✓ |
| FlgN W78A | x | ✓ | ✓ | ✓ |
| FlgN Δ90-100 | x | x | x | x |
| FlgN wild type | ✓ | ✓ | ✓ | ✓ |

### Supplementary Table 4

Table summarising interactions between FliT variants (left column) and export machinery components or cognate subunits (top row). ✓ refers to binding and x refers to weak or no binding.

| **Protein** | FliJ | FlhA | FliD |
|---|---|---|---|
| FliT L72A | x | ✓ | ✓ |
| FliT94 L72A | x | x | ✓ |
| FliT Y106A | ✓ | x | ✓ |
| FliT94 | ✓ | x | ✓ |
| FliT wild type | ✓ | ✓ | ✓ |
